# Functionally and metabolically divergent melanoma-associated macrophages originate from common bone-marrow precursors

**DOI:** 10.1101/2023.06.02.543515

**Authors:** Gabriela A. Pizzurro, Kate Bridges, Xiaodong Jiang, Aurobind Vidyarthi, Kathryn Miller-Jensen, Oscar R. Colegio

## Abstract

Melanomas display high numbers of tumor-associated macrophages (TAMs), which correlate with worse prognosis. Harnessing macrophages for therapeutic purposes has been particularly challenging due to their heterogeneity, based on their ontogeny and function and driven by the tissue-specific niche. In the present study, we used the YUMM1.7 model to better understand melanoma TAM origin and dynamics during tumor progression, with potential therapeutic implications. We identified distinct TAM subsets based on F4/80 expression, with the F4/80^high^ fraction increasing over time and displaying tissue-resident-like phenotype. While skin-resident macrophages showed mixed ontogeny, F4/80^+^ TAM subsets in i.d. YUMM1.7 tumors originated almost exclusively from bone-marrow precursors. Multiparametric analysis of macrophage phenotype showed a temporal divergence of F4/80^+^ TAM subpopulations, which also differed from skin-resident subsets, and from their monocytic precursors. Overall, F4/80^+^ TAMs displayed co-expression of M1- and M2-like canonical markers, while RNA-seq and pathway analysis showed differential immunosuppressive and metabolic profiles. GSEA showed F4/80^high^ TAMs to rely on oxidative phosphorylation, with increased proliferation and protein secretion while F4/80^low^ cells had high pro-inflammatory and intracellular signaling pathways, with lipid and polyamine metabolism. Overall, the present in-depth characterization provides further evidence of the ontogeny of the evolving melanoma TAMs, whose gene expression profiles matched recently-identified TAM clusters in other tumor models and human cancers. These findings provide evidence for potentially targeting specific immunosuppressive TAMs in advanced tumor stages.

## 1. Introduction

Melanoma is the most lethal form of skin cancer. Approximately 200,000 new cases of melanoma will be diagnosed in 2023 in the US, with more than half diagnosed at an invasive stage [1]. Despite being an aggressive disease, melanoma is an immunogenic tumor, and immunotherapy that aims to reprogram the immunosuppressive tumor microenvironment (TME) and boost T cell function has provided a promising approach for melanoma treatment [2,3]. Tumor development represents a special challenge to the host, being a sterile insult yet broadly pathogenic. Several components of the TME are closely related to cancer progression through multiple mechanisms facilitating dissemination and contributing to immune suppression [4].

Macrophages constitute the dominant myeloid cell population in most solid tumors and studies in both humans and mice have linked macrophage density with tumor growth [5]. Tumor-associated macrophages (TAMs) can facilitate tumor progression, proliferation and metastasis by stimulating angiogenesis and inhibiting antitumor T cell responses. Previous results also showed that immunosuppressive TAMs can be functionally reprogrammed to help control tumor growth [3,6]. TAMs were historically considered to be in a ‘M2 state’, referring to the dichotomy of pro-inflammatory (M1) vs anti-inflammatory (M2) cells [5]. Although this classification works well *in vitro*, within a tissue microenvironment, macrophages acquire a wide spectrum of activation states influenced by multiple factors and environmental cues [7]. In an *in vitro* 3D collagen model, we have previously shown that stromal and tumor cells shape macrophage activity in the early melanoma TME. These cell-cell interactions induced bone-marrow (BM)-derived macrophages to acquire an immunosuppressive functional signature that resembled TAMs from melanoma tumors, which differed from the canonical reference polarization states [8].

In recent years, there has been an effort to understand macrophage immune response heterogeneity, from a therapeutic perspective. Expression levels of F4/80 and CD11b during development identified BM-dependent and -independent tissue-colonizing macrophage precursors [9]. The classification of mononuclear phagocytes has also been revisited [10]. Tissue-resident macrophages have been shown to have a distinct origin, gene expression, and phenotype from those derived from CC-chemokine receptor 2 (CCR2)^+^ monocyte precursors recruited to a site of inflammation. The new mononuclear phagocyte classification system, based on cellular origin, allows for a more robust definition across tissues and species [10], and a better understanding of their functional roles during development, homeostasis and disease [11-14]. Under steady-state conditions, BM-derived blood Ly6C^high^ monocytes give rise to two developmental streams: one with CCR2 expression, the acquisition of MHCII molecules, and reduction of Ly6C expression; and the other, tissue-resident macrophages which become Ly6C^-^, CCR2^-^ and CX3CR1^+^, with mixed expression of MHCII molecules [15].

Conditions of tissue stress result in local proliferation of macrophage populations, but monocytes still clearly contribute to the myeloid cell pool under inflammatory conditions, such as during infection or in a growing tumor. Evidence in mouse models such as breast [13], colorectal [16] and pancreatic cancer [17] showed that macrophages in the TME have a mixed embryonic and BM origin, with different levels of expression of lineage markers, proliferation rates and dependence on growth factors, such as CSF1, as previously seen during development [9]. These findings indicate that both monocyte infiltration and proliferation have a role in macrophage maintenance during tumor growth [18].

Another critical factor influencing TAM function is their metabolic state, which impacts immune performance [19]. Glycolysis and mitochondrial oxidative phosphorylation (OXPHOS) are the two principal bioenergetic pathways in a cell, which impact virtually all aspects of cellular metabolism. Cell-selective partitioning of nutrients and metabolic processes in tumor-infiltrating immune cells could be exploited to design novel targeted therapies. Both tumor and tumor-infiltrating immune cells rely on glucose, but it has been shown that cell-intrinsic programs drive preferential acquisition of glucose and glutamine by immune and cancer cells, respectively [20]. A recent study [21] showed that high cellular density in solid tumors can result in lactate build-up in the core. Lactate acts as a quorum-sensing-like signal, leading to increased oscillations in tumor cell HIF-1α activity, rescuing inhibition of cell division and changing gene expression. Our prior findings showed that lactate-driven, HIF-1α-dependent Arginase (Arg)-1 expression in TAMs has an important role in tumor growth [22], possibly via production of polyamines, with a critical role in cell proliferation.

This evidence highlights the need for an in-depth understanding of the origin, evolution and processes that sustain and promote tumor-supportive macrophages. These insights could provide a roadmap for effective therapeutic intervention [23]. In the present report, we focused on profiling multiple factors contributing to macrophage heterogeneity in a fast-growing, aggressive melanoma mouse model with clinically-relevant features, low T-cell infiltration and a dynamic change in the myeloid infiltrate.

## 2. Materials and Methods

### 2.1. Animals

C57BL/6J, B6.129P2-Myb^tm1^Cgn/TbndJ and B6.Cg-Tg^(Mx1-cre)1^Cgn/J mice were purchased from Jackson Laboratories. C57BL/6-Tg^(UBC-GFP)30^Scha/J mice were kindly provided by the Girardi Lab at Yale. Myb^fl/fl^Mx-1^cre/wt^ mice were generated by crossing parental strains. Genotyping was done according to Jax protocols for those strains. Mice were kept according to the standard housing conditions of the Yale Animal Resources Center in specific pathogen-free conditions, and were 8-10 weeks of age at the moment of starting the experiments. All experiments were performed according to the approved protocols of the Yale University Institutional Animal Care and Use Committee.

### 2.2. Cell culture

Yale University Mouse Melanoma (YUMM)1.7 were kindly provided by the Bosenberg Lab at Yale. YUMM1.7 is a clinically-relevant model since it carries the main human melanoma driver mutations BRAFV600E, Pten^−/−^, and Cdkn2^−/−^ [24]. PDVC57 cell line was kindly provided by the Dr. Llanos-Casanova, CIEMAT, Spain. Cell lines were grown in DMEM/F12 media supplemented with 10% FBS, 1% NEAA, 2 mM L-glutamine, 1% Pen/Strep and 1% sodium pyruvate (Gibco).

### 2.3. Tumor studies and sample processing

Mice were anesthetized with isoflurane and intradermally injected with 0.5×10^6^ tumor cells in both flanks, and monitored according to IACUC approved mouse protocol. Tumor volume was calculated as 0.52 x length x width^2^. At experimental endpoint, mice were euthanized in a CO_2_ chamber and tumors were resected and weighted before processing. Briefly, tumors were first cut and then chopped with razor into 1mm^3^ pieces. They were incubated in digestion buffer (1X PBS Ca+Mg+ containing 0.1mg/ml DNase I, Roche 05401127001, and 0.82mg/ml Collagenase IV, from C. histolyticum, Sigma-Aldrich C1889) at 37°C in orbital shaker for 30 min. Samples were vortexed, kept on ice and filtered in 40µm cell strainer. Cells were resuspended in ACK lysis buffer at RT for 5 min. Samples were finally washed, resuspended and counted for downstream processing.

### 2.4 Histology and immunofluorescence

For hematoxylin/eosin (H&E) staining, tumor samples were fixed in formaldehyde and paraffin-embedded at Yale Pathology core, according to standard protocols. The H&E slides were visualized with light microscopy.

For immunofluorescence, freshly collected tumor samples were fixed in 2% paraformaldehyde ON, washed and transferred to 30% sucrose in PBS for 24h, and finally placed in a plastic mold in optimal cutting temperature compound (OCT) at -80°C. Nine □m sections, cut in a Leica cryostat, were washed in PBS and stained with 1:1000 Rb anti-GFP (ab6556, Abcam) and 1:75 Rat anti-mouse F4/80 AF647 (123122, BioLegend) overnight at 4°C. Next, samples were washed and stained with 1:400 Goat anti-Rabbit AF488 and counterstained with DAPI, and analyzed in a Leica SP5 confocal microscope at Yale CCMI Imaging Core.

### 2.5. Flow cytometry

For flow cytometry, we used the following antibodies and dyes (clone, catalog#): CD45 PerCP (30-F11, 103132), CD45.1 APC-Cy7(A20, 110716), CD45.2 APC (104, 109814), B220 FITC (RA3-6B2, 103206), CD3e FITC (KT3.1.1, 155604), NK1.1 FITC (PK136, 108706), CD11b BV421 (M1/70, 101236), CD11b PECy7 (M1/70, 101216), F4/80 PerCP (BM8, 123126) and F4/80 AF700 (BM8, 123130), CD11c PECy7 (N418, 117318),, CD207 PE (4C7, 144204), CSF1R BV605 (AFS98, 135517), Ly6C APC (HK1.4, 128016), Ly6C Pacific Blue (HK1.4, 128014), Ly6G AF647 (1A8, 127610), Ly6G FITC (1A8, 127606), CX3CR1 BV605 (SA011F11, 149027), Arg1 APC (A1exF5, 12-3697-82), CD40 PE (3/23, 124609), CD86 PE-Dazzle594 (GL1,105042), CD64 PE-Dazzle594 (X54-5/7.1, 139320), CD24 BV605 (M1/69, 101827), CD103 BV711 (2E7, 121435), MHCII APC-Cy7 (M5/114.15.2, 107628), CD206 APC (C068C2, 141708) from Biolegend, Live/Dead eFluor506 (65-0866-18), iNOS AF488 (CXNFT, 53-5920- 82), iNOS APC (CXNFT, 17-5920-82) and iNOS PE (CXNFT, 12-5920-82) from Thermo Fisher, and CCR2 PE (475301, FAB5538P-100), Arg1 APC (658922, IC5868A) from R&D Systems. For phenotype analysis, single-cell suspensions were stained in FACS Buffer (PBS 2% FBS). Briefly, cells were incubated with FcBlock 1:200 (anti-CD16/CD32, eBiosciences), washed and stained for extracellular markers. CytoFix/CytoPerm and Perm/Wash Buffer kit (BD, 554714) was used for intracellular staining steps, according to manufacturer’s instructions, and stained for intracellular antigens. Samples analyzed on a LSRFortessa (BD Biosciences). Gating for analysis was performed as described in Supplementary Fig.1a-b and 2a, 2f. Flow cytometry data was analyzed using FlowJo software (TreeStar Inc.). TAM phenotype was further analyzed by principal component analysis (PCA) using SIMCA16 (Sartorius) and GraphPad Prism.

### 2.6. Bone Marrow transplant

Myb^fl/fl^Mx-1^cre/wt^ were administered 10□g/g of Poly(I:C) (P1530, Sigma-Aldrich, USA) i.p. every other day for a total of 7 times, to induce the genetic depletion of BM precursors. C57BL/6-Tg^(UBC-GFP)30^Scha/J mice were treated with 200 μg/g Busulfan (cat#14843, Cayman Chemical, USA) i.p., every day for a total of 5 times, to induce chemical ablation of BM precursors. The same day of the adoptive transfer, BM cells were extracted from the femur and tibia of C57Bl/6J or B6-GFP donor mice (CD45.2^+^), as previously described [25], and 10×10^6^ BM cells were transferred retro-orbitally. Blood samples were extracted from transplanted mice to evaluate chimerism (Myb every 4 weeks, B6-GFP every week). YUMM1.7 tumors were injected after the transplant was stable (Myb after 12 weeks, B6-GFP after 8 weeks), and tumors were processed after 2 weeks, along with blood and skin samples.

### 2.7. Bulk RNA sequencing (RNA-seq) analysis

For TAM sorting, single-cell suspensions from day 21 tumors were prepared as described above, and stained without fixing. This timepoint allowed us to sort the necessary cell numbers of each TAM subpopulation for sequencing. TAMs were sorted in a FACSAria instrument (BD Biosciences) from singlets, Live^+^CD45^+^Lin^-^CD11b^+^ cells and F4/80^high^ and F4/80^low^ were collected and fixed for bulk RNA-seq. Samples were sent to the Yale Center for Genome Analysis (YCGA) for RNA extraction and sequencing. The bulk RNA-seq data was processed with YCGA’s standard alignment pipeline. Protein coding genes with at least ten counts across samples were retained for downstream analyses. The DESeq2 package in R [26] was then used to correct for batch effects, calculate differentially expressed genes (DEGs; adjusted p-value < 0.05 and |log_2_FC| > 1) across experimental conditions, and embed the samples in principal component analysis (PCA) space. The results were visualized using the ggplot2 package in R [27]. Gene set enrichment analysis on the DEGs was performed using the Generally Applicable Gene-set Enrichment (GAGE) package in R [28]. Gene Ontology (GO) biological processes and the Kyoto Encyclopedia of Genes and Genomes (KEGG) were used as reference databases. We performed gene-set enrichment analysis (GSEA) with the online software from UCSD/Broad Institute [29].

### 2.8. Fluorescent bead-based multiplex protein secretion profiling

For supernatant collection, sorted macrophages were plated at a density of 1×10^6^/ml in petri dishes, incubated at 37°C ON and collected, centrifuged and kept at -80°C or directly submitted to Eve Technologies Corp (Calgary, Alberta, Canada). Samples were analyzed with the Mouse Cytokine/Chemokine 44-Plex Discovery Assay® Array (MD44). All detected analytes were within the dynamic range of the standard curves of each analyte, observing no saturation in the samples analyzed. For data analysis purposes, those presenting an out of range (OOR) measurement below the parameter logistic standard curve were systematically replaced with the lowest value obtained for a particular analyte, as per suggestion of the company. The data were natural log-transformed to aid visualization. Samples were visualized and hierarchically clustered using the clustermap function from the Seaborn module in Python. The non-normalized data was embedded in two dimensions using PCA, as implemented in the multivariate statsmodels module in Python.

### 2.9. Statistical analysis

For group comparisons, one-way or two-way ANOVA, with post-hoc comparisons, and linear regressions were performed in GraphPad Prism 9. Significance annotation: ns= not significant, *p<0.05, **p<0.01, ***p<0.001, ****p<0.0001.

## 3. Results

### 3.1. F4/80 expression defined YUMM1.7 melanoma TAM subsets with partial similarities with skin-resident macrophages that evolved during tumor progression

YUMM1.7 tumors show an overall low immune infiltration [30]. In YUMM1.7 i.d. tumors evaluated 14 days post-injection, we detected a majority of myeloid infiltrate, with macrophages being the predominant myeloid cell type (Fig. 1a). YUMM1.7 cells generated fast-growing i.d. tumors in C57Bl/6 mice, which infiltrated the whole skin compartment, colonizing the dermis and displaying an inflamed epidermis (Supplementary Fig. 1c-d). When looking at Lin^-^MHCII^+^ cells, only a small fraction of them corresponded to dendritic cells (cDC1 and cDC2) or Langerhans cells, confirming TAM prevalence (Supplementary Fig. 1e). CD11b^+^F4/80^+^ macrophages were found in both YUMM1.7 tumors and normal skin (Fig. 1b). Looking closer into F4/80^+^ TAMs, there was a clear separation between F4/80^low^- and F4/80^high^-expressing cells, where F4/80^high^ TAMs represented a smaller fraction of the total, compared to F4/80^low^ and F4/80^neg^ TAMs (Fig. 1c-d). The macrophage F4/80-based subset distribution was mirrored the in the skin. Although TAMs were found throughout the tumors, showing enriched regions with F4/80^+^ clusters, macrophages were only sparse in the skin compartment (Supplementary Fig. 1f-g).

**Figure 1.**
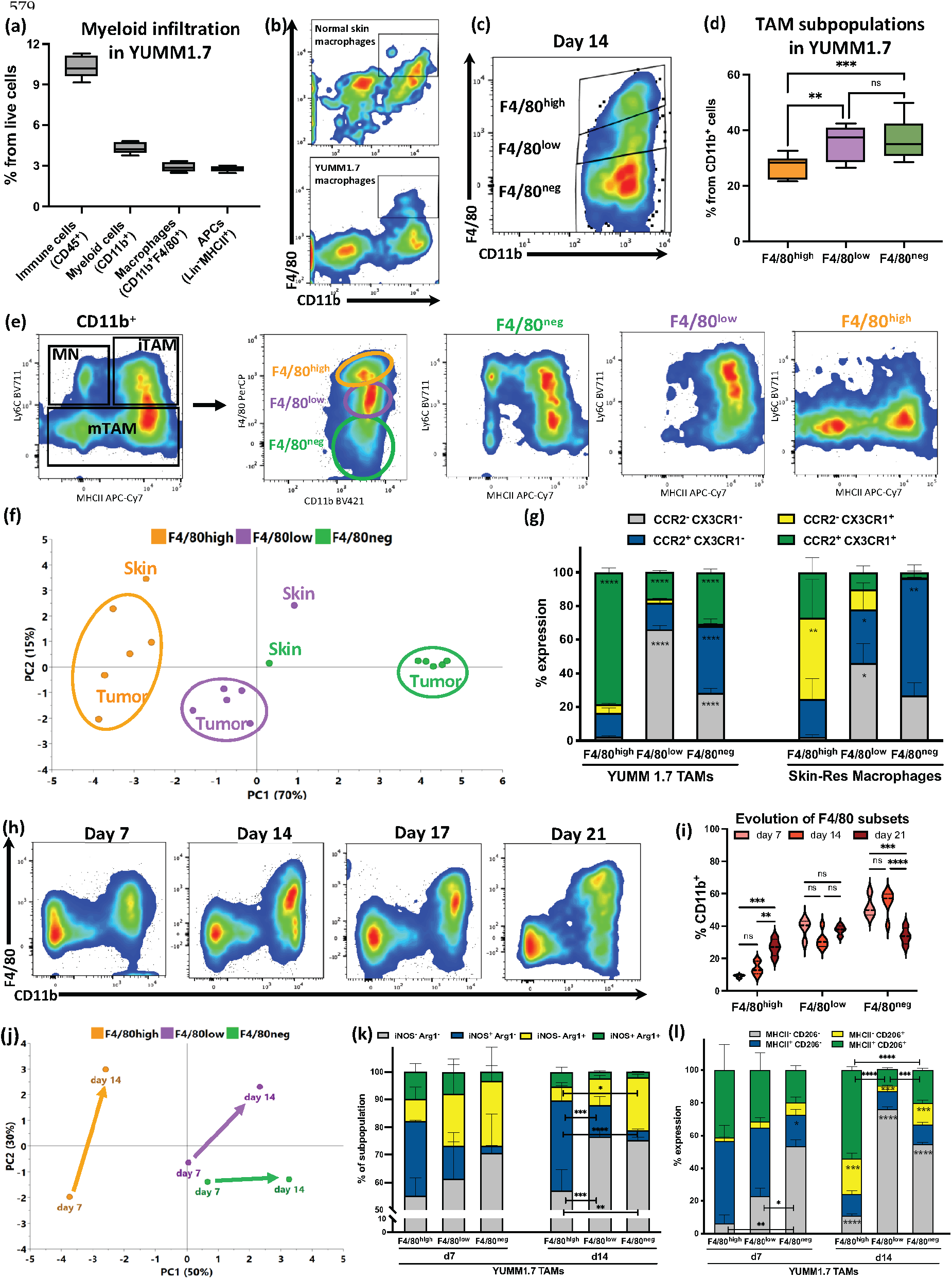
Characterization and evolution of F4/80^+^ tumor-associated macrophage subsets in intradermal melanoma tumors. **(a)** Box plots of FACS quantification of YUMM1.7 immune infiltration. Percentage of total immune infiltration (CD45^+^), myeloid cells (CD45^+^CD11b^+^) and APC (CD45^+^Lin^-^MHCII^+^). n=10, pooled from at least 3 independent experiments. **(b)** Representative flow cytometry plots gating the macrophages (CD11b^+^F4/80^+^) out of the total CD45^+^ cells, both in normal skin and in YUMM1.7 i.d. tumors. **(c)** Representative plot of the Lin^-^CD11b^+^ myeloid subpopulations determined by their F4/80 expression in YUMM1.7 melanoma tumors. **(d)** Box plots with FACS quantification of F4/80^+/-^ myeloid subpopulations at day 14 YUMM1.7 i.d. tumors. n=15, pooled from 3 independent experiments. **(e)** Representative flow cytometry plots of the expression of Ly6C and MHCII markers (‘monocyte waterfall’), defining myeloid cell subsets, broken down into the different F4/80^+/-^ subpopulations in YUMM1.7 tumors. MN=monocytes; iTAM=immature TAMs; mTAMs=mature TAMs. **(f)** PCA plot of the tumor-infiltrating and skin-resident F4/80 subpopulations phenotype, assessed by flow cytometry. Details of flow panels and parameters included in the PCA can be found in Supp. Fig 1h-j. 5 independent tumor samples from day 14 were included in the analysis. Skin macrophage phenotype data is plotted as reference subpopulations, average of 2 independent samples. **(g)** Expression of chemokine receptors CCR2/CX3CR1 in F4/80 macrophage subsets in YUMM1.7 tumors, compared to skin-resident macrophage subsets. Statistical analysis showing comparisons between marker-expressing subsets within each macrophage type. TAMs n=5 and Skin n=3, pooled from at least 2 independent experiments. **(h)** Representative flow cytometry plots showing the changes of TAM subpopulations over time in YUMM1.7 i.d. tumors. **(i)** Quantification of the evolution of the F4/80 TAM subpopulations, assessed by flow cytometry, at days 7, 14 and 21 after YUMM1.7 i.d. tumor injection. n=4-6, pooled from independent experiments. **(j)** PCA plot of tumor-infiltrating F4/80 subpopulations phenotype at day 7 and day 14 YUMM1.7 i.d. tumors in B6 mice. Details of flow panels and parameters included in the PCA can be found in Supp. Fig 1h, l-m. For day 7 tumors, n=3 and day 14 tumors, n=5. **(k)** and **(l)** Analysis of canonical M1/M2-marker pairs co-expression (iNOS/Arginase-1 and MHCII/CD206) in TAM subpopulations from YUMM1.7 tumors, at day 7 and 14. Statistical analysis showing comparisons between marker-expressing subsets within each timepoint. n=3-5 pooled mice per group, from at least 2 independent experiments. ns= not significant, *p<0.05, **p<0.01, ***p<0.001, ****p<0.0001.

When mapped into the monocyte classification based on Ly6C and MHCII expression [31,32], the F4/80^high^ subpopulation resembled skin tissue-resident macrophages, while the other subsets aligned with monocyte-derived cells (Fig. 1e). We had previously determined that YUMM melanoma TAMs share features with both canonical *in vitro* M1-like and M2-like profiles, and that BMDMs can be shaped into a TAM-like state through cell-cell communications and interactions in a 3D environment [8]. To study melanoma TAMs in more detail, we phenotypically profiled macrophages from the tumor and normal skin by flow cytometry (Supplementary Fig. 1h-i). We used PCA to compare these multi-dimensional phenotypic profiles. PC1 separated samples by F4/80 expression level, which explained most of the variance in these data, and clearly partitioned TAM subsets (Fig. 1f, Supplementary Fig. 1j). F4/80^high^ TAMs were the most heterogeneous between samples and the more similar to the matched F4/80 skin subsets (Fig. 1f).

We examined individual marker expression between tumor- vs skin-infiltrating macrophages. Skin-resident macrophage chemokine receptor expression patterns showed F4/80 expression to be inversely correlated to CCR2 expression and positively related to expression of CX3CR1 (Fig. 1g). Although TAM subsets had differences in their chemokine receptor expression, they did not follow the same pattern as skin-resident cells, with an overall higher co-expression of both markers (Fig. 1g). Similarly, skin monocyte-derived cells had decreasing levels of CSF1R expression as they became more ‘mature’ cells (i.e. MHCII^+^ F4/80^high^), while TAMs retained significant levels of CSF1R expression, and MHCII^-^ cells as F4/80 expression increased (Supplementary Fig. 1k).

We then looked into how F4/80 TAM subsets in the YUMM1.7 tumors changed over time. Interestingly, F4/80^+^ TAM fraction was enriched from day 7 to day 21, primarily due to a significant increase in the F4/80^high^ subpopulation (Fig. 1h-i). TAM subsets also exhibited a divergence in their phenotype as the tumor progressed (Fig. 1j, Supplementary Fig. 1l). When we looked into canonical M1/M2 markers, there was no clear association between our TAM subsets and their evolution with reference to macrophage polarization states, as we have previously described for the general YUMM1.7 TAM population [8]. Higher expression of iNOS and MHCII were present in more immunosuppressive subsets and advanced timepoints (Fig. 1k-l). Of note, a higher percentage of expression of iNOS did not necessarily correlate with a higher mean fluorescent intensity, potentially impacting the overall immune performance (Supplementary Fig. 1l).

### 3.2. Despite exhibiting some skin-resident-like features, melanoma TAMs originated almost exclusively from circulating monocytes

Given the partial similarities between melanoma TAMs and skin-resident cells and observations in other tumor types [13,16,17,33], we further investigated TAM ontogeny to better understand their immune function. For this, we used two strategies to track BM precursors and tumor infiltration (Fig. 2a, Supplementary Fig. 2a). Chemical ablation with busulfan model provided an almost complete and homogenous BM transplant, while genetic depletion of Myb^+^ BM precursor generated a graded engraftment success, allowing for evaluation of potential competition between host and donor cells (Fig. 2b, Supplementary Fig. 2b). The busulfan-treated mice showed significantly better myeloid cell engraftment, observed in circulation, which allowed us to evaluate the F4/80 TAM origin. At day 14, all YUMM1.7 TAMs originated from GFP^-^ donor BM precursors (Fig. 2c, Supplementary Fig. 2c).

**Figure 2.**
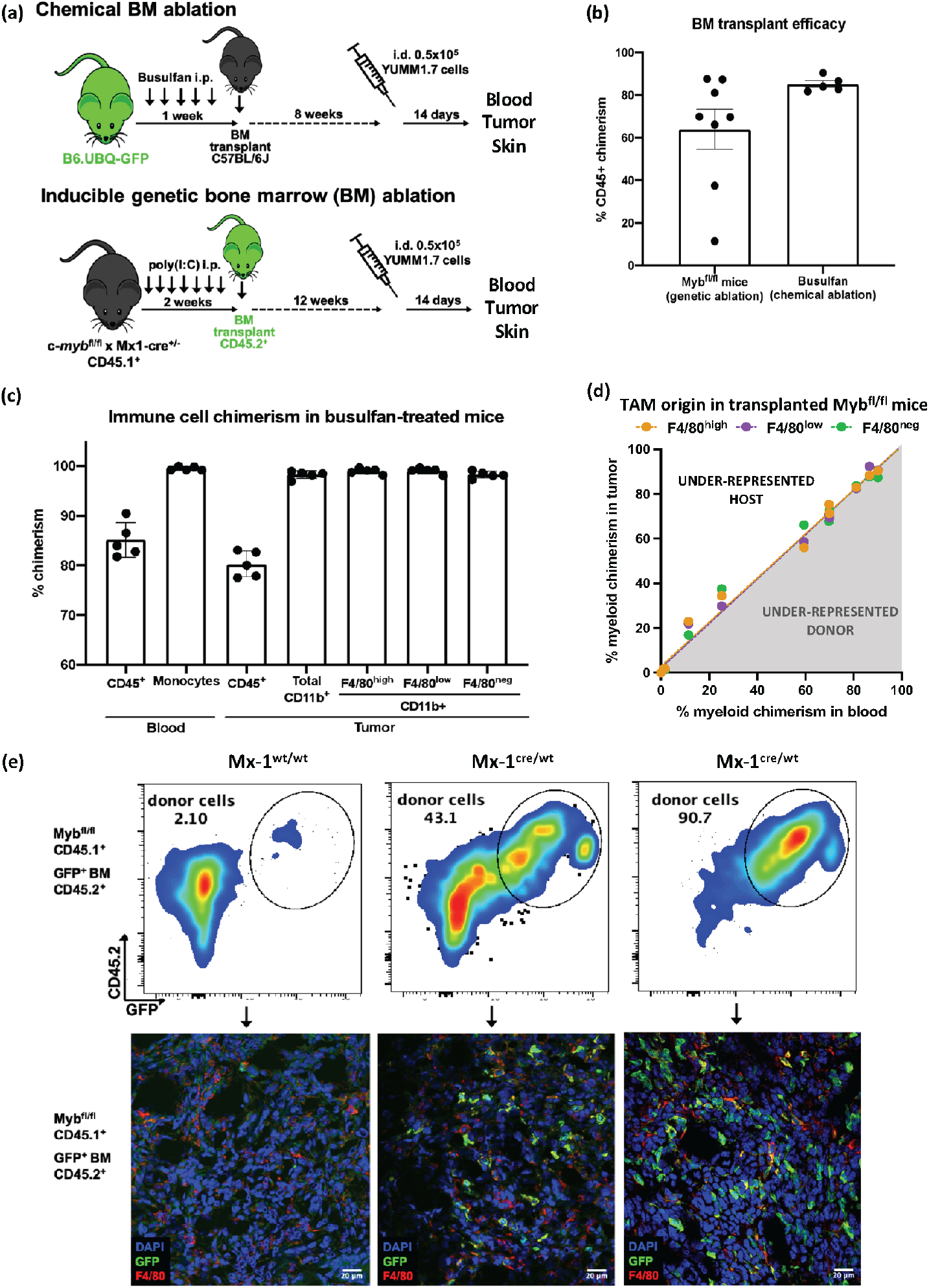
Determination of the origin of tumor-infiltrating myeloid cells in YUMM1.7 melanoma tumors. **(a)** Schematic design showing two different strategies used to perform bone-marrow transplants and track the origin of YUMM1.7-infiltrating macrophages. **(b)** BM transplant efficacy shown as percentage of chimerism of CD45^+^ cells in transplanted mice blood. Chimerism is expressed as the percentage of donor cells in the host, in both BM transplant approaches. **(c)** Tumor-infiltrating myeloid cell origin in YUMM1.7 tumors in busulfan-treated mice. Percentages of chimerism of total CD45^+^ cells and myeloid cells were compared between blood and tumor compartments. n=5. **(d)** Assessment of macrophage origin in YUMM1.7 tumors in Myb^fl/fl^Mx-1^cre/wt^ and Myb^fl/fl^Mx-1^wt/wt^ mice. Linear regressions were performed to compare the chimerism of myeloid cells observed in circulation and the chimerism of myeloid tumor-infiltrating cells. TAM subsets were analyzed separately. Tumor samples, n=12, pooled from 3 independent experiments. **(e)** In Myb^fl/fl^Mx-1^cre/wt^ mice with increasing levels of BM donor (GFP^+^, CD45.2^+^) engraftment, macrophage infiltration and chimerism was validated by immunofluorescence on respective tumor slides. Donor cells were detected by GFP staining and TAMs by F4/80^+^.

In the Myb^fl/fl^Mx-1^cre^ model, YUMM1.7 showed similar tumor growth and all F4/80 subsets were observed in the infiltrate (Supplementary Fig. 2d-e). In the genetic ablation model, we again used the chimerism observed in circulating cells to analyze the tumor and skin myeloid infiltration (Supplementary Fig. 2f). Once again, the evidence showed that all melanoma TAMs derived from BM precursors, since chimerism of each subset within the tumor matched the host/donor monocyte proportion in circulation (Fig. 2d). We were able to validate this approach in the mouse skin (Supplementary Fig. 2g). F4/80^high^ macrophages, previously described to be Myb-independent during development and to colonize the tissue and self-renew [9], showed lower levels of chimerism in the skin compared to monocytes in circulation (Supplementary Fig. 2h). We also analysed TAMs in a non-melanoma PDVC57 skin cancer model [34] to compare the macrophage infiltration in the dermal compartment (Supplementary Fig. 2 i-j). In the PDVC57 tumors, we observed the F4/80-based TAM subsets as seen in YUMM1.7 tumors. But, interestingly, while F4/80^low^ TAMs were exclusively originated from BM precursors, F4/80^high^ TAMs showed a mixed origin. Based on these observations, the TAM origin in YUMM1.7 could not be exclusively attributed to the dermal location of the tumors. We further validated the macrophage infiltration via tissue immunofluorescence. As seen in Fig. 2e, increasing levels of chimerism corresponded to higher GFP^+^ F4/80^+^ in the tumor, while Mx-1^wt/wt^ controls with no engraftment showed no GFP^+^ cells.

In the busulfan-treated mice, with over 95% of myeloid donor cells, only non-immune stromal cells were GFP^+^ (Supplementary Fig. 2i). Authors should discuss the results and how they can be interpreted from the perspective of previous studies and of the working hypotheses. The findings and their implications should be discussed in the broadest context possible. Future research directions may also be highlighted.

### 3.3. Melanoma F4/80^+^ TAM subsets have distinct immunosuppressive profiles with specific metabolic and functional pathways

Despite a common BM origin, F4/80^+^ melanoma TAMs evolved into phenotypically diverging subsets. To understand the differences between TAM subsets and how those could be leveraged for potential therapeutic strategies, we performed bulk RNA-seq. We collected data from F4/80^high^ and F4/80^low^ TAM subsets sorted from YUMM1.7 tumors in three independent experiments, with approximately 0.15-1.50×10^6^ cells per replicate (Supplementary Fig. 3a). Considering only protein coding genes, F4/80^high^ and F4/80^low^ samples separated in a two-dimensional space by PCA and unsupervised hierarchical clustering revealed distinct transcriptional patterns across TAM subsets (Fig. 3a, Supplementary Fig. 3b). Identification of the differentially expressed genes (DEGs) revealed key transcriptional differences between the F4/80^high^ and F4/80^low^ melanoma TAMs (Supp. Fig. 3c, top 50). Interestingly, both TAM subsets expressed mutually exclusive groups of M2-like, immunosuppressive markers. Of note, F4/80^low^ TAMs overexpressed *Chil3, Mmp9* and *Ear2*, while F4/80^high^ TAMs had elevated expression of scavenger receptors *Mrc1, Mertk* and *Cd163* (Fig. 3b). Each TAM subset also showed upregulation of specific chemokines and chemokine receptors. Most importantly, these two TAM subpopulations have gene profiles similar to TAM clusters previously identified in other tumor models, and in the human setting [23,35-37].

**Figure 3.**
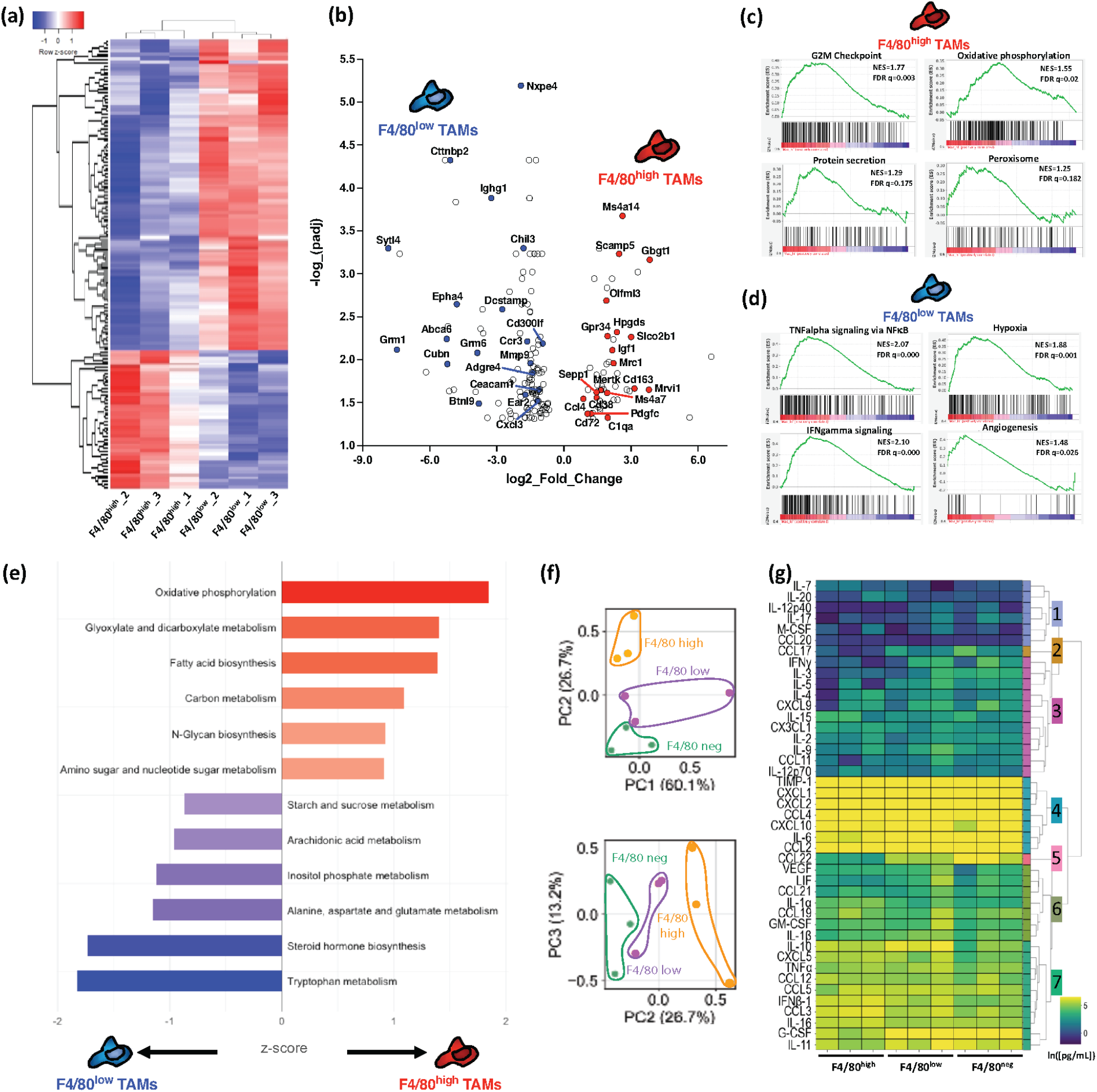
Functional characterization and profiling of TAM subsets in YUMM1.7 melanoma tumors. **(a)** Heatmap with hierarchical clustering of TAM subpopulations based on z-score-transformed gene expression from bulk RNA-seq. F4/80^high^ and F4/80^low^ subsets were isolated from 10 pooled tumors, and RNA-seq data was collected in 3 independent experiments. **(b)** Volcano plot showing the top 50 differentially-expressed genes (DEGs) between F4/80^high^ and F4/80^low^ subsets (p-adj < 0.05, abs(log2FoldChange(FC)) > 1). **(c)** and **(d)** GSEA Hallmark gene sets from the mouse MSigDB, with an FDR<25% for the F4/80^high^ and F4/80^low^ TAM DEGs (NES= normalized enrichment score, FDR= false discovery rate). **(e)** Enrichment of F4/80^high^ (right, red) and F4/80^low^ (left, blue) DEGs (protein coding genes only) for KEGG metabolic pathway terms. Bar lengths represent associated z-score. **(f)** PCA embedding of TAM samples from cytokine/chemokine functional profiling, assessed by bead-based multiplex secretion assay (PC2 vs. PC1, top; PC3 vs. PC2, bottom). Samples clearly separated by TAM subset along PC2, which explained 26.7% of variability in the data. **(g)** Heatmap of natural log-transformed protein secretion from (F). Hierarchical clustering identified 7 functional clusters across TAM subsets.

We further identified clear differences between the cellular and metabolic programs of F4/80^high^ and F4/80^low^ TAM subsets. Gene-set enrichment analysis (GSEA) (Supplementary Fig. 3d) revealed that F4/80^high^ TAMs were enriched for oxidative phosphorylation, with expression of *Igf1* and *Cd38*, and lipid metabolism through the peroxisome pathway, while also showing increased proliferation pathways, such as the G2M checkpoint, and upregulated protein secretion (Fig. 3b-c, 3e, Supplementary Fig. 3d-e). In contrast, F4/80^low^ cells were enriched for proinflammatory signaling pathways, such as TNFα signaling via NFκB, IFNα and IFNγ signaling, along with responses to hypoxia and angiogenesis. These also showed expression of glycolysis-associated ABC- and glutamate-transporter genes, along with upregulated glycolysis KEGG metabolic pathways (Fig. 3d-e, Supplementary Fig. 3e). These results highlighted important differences in terms of metabolic heterogeneity and proliferative capacity and self-renewal of the YUMM1.7 melanoma TAMs.

Functional protein secretion showed differences in the melanoma F4/80 TAM subsets at the protein levels in growth factors, pro-inflammatory and immunosuppressive cytokines, and immune trafficking chemokines. These results were obtained from profiling F4/80^high^, F4/80^low^ and F4/80^neg^ TAM subsets sorted from YUMM1.7 tumors through a multiplex bead-based secretion assay. To validate the functional differences and pathways inferred from RNA-seq analysis we screened for cytokines and chemokines with relevant immune functions in the TME. The PCA-embedded secretion data showed that, despite heterogeneity across replicates, the TAM subsets separated along PC2 (Fig. 3f), with top coefficients leading to factors with differential secretion across TAM subsets (Supp. Fig. 3f). To identify the differences driving functional divergence between F4/80 subsets, we looked into the secreted factors individually (Supp. Fig. 3g). When contrasting F4/80^high^ and F4/80^low^ TAMs, F4/80^high^ showed significant higher levels of IL-15, IL-16 and CX3CL1, and higher trends for IFNβ1, IL-1α, CCL3, CXCL5, IL-20, M-CSF and IL-4, while F4/80^low^ secreted more VEGF, GM-CSF, IL-1β, IL-10, CCL2, CXCL1, CXCL2, CXCL10 and CCL22, with trends for G-CSF, TNF-α, IFN-γ, IL-5, IL-9 and CCL11 (Supp. Fig. 3g). These profiles partially validated some of the transcriptome profiles, adding to the phenotypic differences established by flow cytometry. Finally, we hierarchically clustered the bulk secretion data to define patterns in the trends contributing to differences in F4/80 TAM subsets. We identified 7 functional clusters, determining overall low or high secretion across subsets (clusters 1 and 4), a clear trend across the subsets (clusters 2 and 5), while the others resulted harder to interpret primarily due to sample-to-sample heterogeneity (Fig. 3g).

## 4. Discussion

Previous seminal work [9] had shown that F4/80^bright^ and yolk-sac macrophages had increased expression of receptors *Cx3cr1* and *Csf1*, along with proliferation markers. In particular, the YUMM1.7 F4/80^high^ TAMs displayed similarities with skin-resident macrophages. However, key phenotypic markers, such as CSF1R and chemokine receptors, showed strong differences, which made them resemble to recruited monocytes. Interestingly, fate-mapping of monocytes determined that all melanoma TAMs had a BM origin, although they developed into two separate subsets with different phenotypic profiles. Transcriptional profiling showed F4/80^high^ subset to have features resembling tissue-resident macrophages from an embryonic origin, such as microglia. This phenomenon has been seen in other tissue contexts [38], emphasizing the role of the specific environment for shaping macrophage identity.

Several groups have studied macrophage ontogeny in other tumor types, but mostly in orthotopic settings [13,16,17,37]. In the dermal compartment, we observed different ontogeny in the macrophage composition in i.d. skin tumors. Whether F4/80^+^ TAM subsets and evolution are driven and determined by the skin microenvironment or if it is encoded by each specific TME type will require further studies. However, recent spatial mapping of immune landscape in tumors have shown that completely different functional immune environments can be few millimeters away [39], and that the specific location in the TME, and the location of the tumor itself, influences immune phenotype and function [33,40,41]. We observed differences in TAM distribution by region throughout the YUMM1.7 tumors. A thorough mapping of melanoma TAMs within the TME, through the markers we and others identified in each F4/80 subset, would help to determine immune neighborhoods, understand their cell-cell interactions and evolution during tumor progression, and assess novel targeted therapies, in order to improve future treatment rationale [42-44].

A prior study from our group with a genetically-engineered mouse model (GEMM), which in turn originated the YUMM cell lines, showed differences in F4/80 expression in TAMs, which secreted Chi3l3, MMP9 and IGF1, among other factors [3]. Specific myeloid-targeting therapies CD40-agonist and CSF1R inhibitor changed the F4/80^+^ TAM composition and induced tumor control. The functional divergence of these F4/80^+^ TAMs, with distinct active metabolic pathways, opens another road for thinking therapeutic approaches [19,20,45]. Tissue-resident-like TAMs, which have a longer lifespan, the potential to self-renew and expand through proliferation, could be labeled as the ‘bad guys’ in the TME, which may be more difficult to be externally reprogramed into tumoricidal cells with traditional immunotherapies. A metabolic reprogramming approach gains more importance in this context since it could target cell-intrinsic programs which could impact directly on immune function [46,47]. Interestingly, our RNA-seq signatures matched TAM clusters that were recently-identified through scRNAseq gene signatures from different tumor types and pathologies [23,35,36,48,49]. The skin compartment may provide an ideal setting to test novel approaches [50], since topical application may prevent an otherwise high systemic toxicity.

## 5. Conclusions

The results presented in this report provide an in-depth characterization of the evolution of melanoma TAMs, in a mouse model that resembles the human disease. Unlike other previously studied orthotopic tumor models, such as lung or pancreatic cancer, i.d. melanomas exhibited primarily BM-derived TAMs. More interestingly, they presented in discrete, phenotypically and functionally distinct subsets, with evolving ratios. We comprehensively analyzed melanoma TAM ontogeny and how discrete TAM subsets, defined by their F4/80 expression, show distinct cellular and functional profiles from early stages, and continue to evolve throughout tumor progression.

The TAM similarities between tumor models might help to infer about TAM evolution in the TME and their heterogeneity at the single-cell level. More importantly, we could speculate about potential treatment options. A major challenge as we look forward, though, is centered in how to translate the investigation on macrophage origins, differentiation, and maintenance into humans. Fortunately, in recent years have been advances in several tumor types trying to validate the discoveries made with mouse models [23,33,37,40]. Ultimately, we want to have a better understanding of the heterogeneity and functional characteristics of human TAMs to improve prognosis and treatment options.

## Supplementary Materials

The following supporting information can be downloaded at: www.mdpi.com/xxx/s1, Figure S1: Tumor-associated macrophage subsets in the i.d.-injected YUMM mouse melanoma model; Figure S2: Origin of melanoma-associated macrophage subsets; Figure S3: Gene expression profiling and functional analysis of TAM subsets in YUMM1.7 tumors.

## Author Contributions

Conceptualization, G.A.P. and O.R.C.; methodology, G.A.P; formal analysis, G.A.P. and K.B.; investigation, G.A.P., X.J. and A.V.; resources, O.R.C. and K.M.J.; data curation, K.B.; writing—original draft preparation, G.A.P., K.B. and K.M.J; visualization, G.A.P. and K.B.; supervision, O.R.C. and K.M.J; funding acquisition, O.R.C and K.M.J. All authors have read and agreed to the published version of the manuscript.

## Institutional Review Board Statement

The animal study protocol was approved by the Yale University’s Institutional Animal Care and Use Committee (protocols #2018-20039, from 05/25/2018 to 04/30/2021 and #2019-20085, from 2/13/2019 to 01/31/2022).

## Data Availability Statement

The datasets generated and analyzed in this study are publicly available and can be found in the GEO database:

https://www.ncbi.nlm.nih.gov/geo/query/acc.cgi?acc=GSE223545

## Acknowledgments

We would like to thank the Yale Flow Cytometry, Confocal Imaging and YCGA Cores and technicians at the Department of Pathology at Yale for their assistance and processing of tumor samples. We thank Dr Nanda Forni, and all past and current members of the Colegio and Miller-Jensen lab for helpful discussions. Also, we thank Erin Tracy for her role as research liaison between our labs, and particularly to Dr Gyorgy Paragh, Chair of Dermatology at Roswell Park Comprehensive Cancer Center, for his guidance and support during the manuscript preparation, to honor Dr Oscar Colegio’s work, legacy and memory.

## References

1. Siegel, R.L.; Miller, K.D.; Wagle, N.S.; Jemal, A. Cancer statistics, 2023. CA Cancer J Clin 2023, 73, 17–48, doi:10.3322/caac.21763.

2. Mantovani, A.; Allavena, P.; Marchesi, F.; Garlanda, C. Macrophages as tools and targets in cancer therapy. Nat Rev Drug Discov 2022, 21, 799–820, doi:10.1038/s41573-022-00520-5.

3. Perry, C.J.; Munoz-Rojas, A.R.; Meeth, K.M.; Kellman, L.N.; Amezquita, R.A.; Thakral, D.; Du, V.Y.; Wang, J.X.; Damsky, W.; Kuhlmann, A.L.; et al. Myeloid-targeted immunotherapies act in synergy to induce inflammation and antitumor immunity. J Exp Med 2018, 215, 877–893, doi:10.1084/jem.20171435.

4. Hu, Q.; Wu, G.; Wang, R.; Ma, H.; Zhang, Z.; Xue, Q. Cutting edges and therapeutic opportunities on tumorassociated macrophages in lung cancer. Front Immunol 2022, 13, 1007812, doi:10.3389/fimmu.2022.1007812.

5. Sica, A.; Mantovani, A. Macrophage plasticity and polarization: in vivo veritas. J Clin Invest 2012, 122, 787–795, doi:10.1172/JCI59643.

6. Hoves, S.; Ooi, C.H.; Wolter, C.; Sade, H.; Bissinger, S.; Schmittnaegel, M.; Ast, O.; Giusti, A.M.; Wartha, K.; Runza, V.; et al. Rapid activation of tumor-associated macrophages boosts preexisting tumor immunity. J Exp Med 2018, 215, 859–876, doi:10.1084/jem.20171440.

7. Murray, P.J.; Allen, J.E.; Biswas, S.K.; Fisher, E.A.; Gilroy, D.W.; Goerdt, S.; Gordon, S.; Hamilton, J.A.; Ivashkiv, L.B.; Lawrence, T.; et al. Macrophage activation and polarization: nomenclature and experimental guidelines. Immunity 2014, 41, 14–20, doi:10.1016/j.immuni.2014.06.008.

8. Pizzurro, G.A.; Liu, C.; Bridges, K.; Alexander, A.F.; Huang, A.; Baskaran, J.P.; Ramseier, J.; Bosenberg, M.W.; Mak, M.; Miller-Jensen, K. 3D Model of the Early Melanoma Microenvironment Captures Macrophage Transition into a Tumor-Promoting Phenotype. Cancers (Basel) 2021, 13, pdoi:10.3390/cancers13184579.

9. Schulz, C.; Gomez Perdiguero, E.; Chorro, L.; Szabo-Rogers, H.; Cagnard, N.; Kierdorf, K.; Prinz, M.; Wu, B.; Jacobsen, S.E.; Pollard, J.W.; et al. A lineage of myeloid cells independent of Myb and hematopoietic stem cells. Science 2012, 336, 86–90, doi:10.1126/science.1219179.

10. Guilliams, M.; Dutertre, C.A.; Scott, C.L.; McGovern, N.; Sichien, D.; Chakarov, S.; Van Gassen, S.; Chen, J.; Poidinger, M.; De Prijck, S.; et al. Unsupervised High-Dimensional Analysis Aligns Dendritic Cells across Tissues and Species. Immunity 2016, 45, 669–684, doi:10.1016/j.immuni.2016.08.015.

11. Gomez Perdiguero, E.; Klapproth, K.; Schulz, C.; Busch, K.; Azzoni, E.; Crozet, L.; Garner, H.; Trouillet, C.; de Bruijn, M.F.; Geissmann, F.; et al. Tissue-resident macrophages originate from yolk-sac-derived erythro-myeloid progenitors. Nature 2015, 518, 547–551, doi:10.1038/nature13989.

12. Hashimoto, D.; Chow, A.; Noizat, C.; Teo, P.; Beasley, M.B.; Leboeuf, M.; Becker, C.D.; See, P.; Price, J.; Lucas, D.; et al. Tissue-resident macrophages self-maintain locally throughout adult life with minimal contribution from circulating monocytes. Immunity 2013, 38, 792–804, doi:10.1016/j.immuni.2013.04.004.

13. Franklin, R.A.; Liao, W.; Sarkar, A.; Kim, M.V.; Bivona, M.R.; Liu, K.; Pamer, E.G.; Li, M.O. The cellular and molecular origin of tumor-associated macrophages. Science 2014, 344, 921–925, doi:10.1126/science.1252510.

14. Gautier, E.L.; Shay, T.; Miller, J.; Greter, M.; Jakubzick, C.; Ivanov, S.; Helft, J.; Chow, A.; Elpek, K.G.; Gordonov, S.; et al. Gene-expression profiles and transcriptional regulatory pathways that underlie the identity and diversity of mouse tissue macrophages. Nat Immunol 2012, 13, 1118–1128, doi:10.1038/ni.2419.

15. Baranska, A.; Shawket, A.; Jouve, M.; Baratin, M.; Malosse, C.; Voluzan, O.; Vu Manh, T.P.; Fiore, F.; Bajenoff, M.; Benaroch, P.; et al. Unveiling skin macrophage dynamics explains both tattoo persistence and strenuous removal. J Exp Med 2018, 215, 1115–1133, doi:10.1084/jem.20171608.

16. Soncin, I.; Sheng, J.; Chen, Q.; Foo, S.; Duan, K.; Lum, J.; Poidinger, M.; Zolezzi, F.; Karjalainen, K.; Ruedl, C. The tumour microenvironment creates a niche for the self-renewal of tumour-promoting macrophages in colon adenoma. Nat Commun 2018, 9, 582, doi:10.1038/s41467-018-02834-8.

17. Zhu, Y.; Herndon, J.M.; Sojka, D.K.; Kim, K.W.; Knolhoff, B.L.; Zuo, C.; Cullinan, D.R.; Luo, J.; Bearden, A.R.; Lavine, K.J.; et al. Tissue-Resident Macrophages in Pancreatic Ductal Adenocarcinoma Originate from Embryonic Hematopoiesis and Promote Tumor Progression. Immunity 2017, 47, 323–338 e326, doi:10.1016/j.immuni.2017.07.014.

18. Franklin, R.A.; Li, M.O. Ontogeny of Tumor-associated Macrophages and Its Implication in Cancer Regulation. Trends Cancer 2016, 2, 20–34, doi:10.1016/j.trecan.2015.11.004.

19. Hong, H.S.; Mbah, N.E.; Shan, M.; Loesel, K.; Lin, L.; Sajjakulnukit, P.; Correa, L.O.; Andren, A.; Lin, J.; Hayashi, ; et al. OXPHOS promotes apoptotic resistance and cellular persistence in T(H)17 cells in the periphery and tumor microenvironment. Sci Immunol 2022, 7, eabm8182, doi:10.1126/sciimmunol.abm8182.

20. Reinfeld, B.I.; Madden, M.Z.; Wolf, M.M.; Chytil, A.; Bader, J.E.; Patterson, A.R.; Sugiura, A.; Cohen, A.S.; Ali, A.; Do, B.T.; et al. Cell-programmed nutrient partitioning in the tumour microenvironment. Nature 2021, 593, 282–288, doi:10.1038/s41586-021-03442-1.

21. Kshitiz; Afzal, J.; Suhail, Y.; Chang, H.; Hubbi, M.E.; Hamidzadeh, A.; Goyal, R.; Liu, Y.; Sun, P.; Nicoli, S.; et al. Lactate-dependent chaperone-mediated autophagy induces oscillatory HIF-1alpha activity promoting proliferation of hypoxic cells. Cell Syst 2022, doi:10.1016/j.cels.2022.11.003.

22. Colegio, O.R.; Chu, N.Q.; Szabo, A.L.; Chu, T.; Rhebergen, A.M.; Jairam, V.; Cyrus, N.; Brokowski, C.E.; Eisenbarth, S.C.; Phillips, G.M.; et al. Functional polarization of tumour-associated macrophages by tumour-derived lactic acid. Nature 2014, 513, 559–563, doi:10.1038/nature13490.

23. Zhang, L.; Li, Z.; Skrzypczynska, K.M.; Fang, Q.; Zhang, W.; O’Brien, S.A.; He, Y.; Wang, L.; Zhang, Q.; Kim, A.; et al. Single-Cell Analyses Inform Mechanisms of Myeloid-Targeted Therapies in Colon Cancer. Cell 2020, 181, 442–459 e429, doi:10.1016/j.cell.2020.03.048.

24. Meeth, K.; Wang, J.X.; Micevic, G.; Damsky, W.; Bosenberg, M.W. The YUMM lines: a series of congenic mouse melanoma cell lines with defined genetic alterations. Pigment Cell Melanoma Res 2016, 29, 590–597, doi:10.1111/pcmr.12498.

25. Trouplin, V.; Boucherit, N.; Gorvel, L.; Conti, F.; Mottola, G.; Ghigo, E. Bone marrow-derived macrophage production.J Vis Exp 2013, e50966, doi:10.3791/50966.

26. Love, M.I.; Huber, W.; Anders, S. Moderated estimation of fold change and dispersion for RNA-seq data with ESeq2. Genome Biol 2014, 15, 550, doi:10.1186/s13059-014-0550-8.

27. Wickham, H. ggplot2 Elegant Graphics for Data Analysis Introduction. Use R 2009, 1-+, doi:10.1007/978-0-387-98141-3_1.

28. Luo, W.; Friedman, M.S.; Shedden, K.; Hankenson, K.D.; Woolf, P.J. GAGE: generally applicable gene set enrichment for pathway analysis. BMC Bioinformatics 2009, 10, 161, doi:10.1186/1471-2105-10-161.

29. Subramanian, A.; Tamayo, P.; Mootha, V.K.; Mukherjee, S.; Ebert, B.L.; Gillette, M.A.; Paulovich, A.; Pomeroy, S.L.; Golub, T.R.; Lander, E.S.; et al. Gene set enrichment analysis: a knowledge-based approach for interpreting genomewide expression profiles. Proc Natl Acad Sci U S A 2005, 102, 15545–15550, doi:10.1073/pnas.0506580102.

30. Wang, J.; Perry, C.J.; Meeth, K.; Thakral, D.; Damsky, W.; Micevic, G.; Kaech, S.; Blenman, K.; Bosenberg, M. UV-induced somatic mutations elicit a functional T cell response in the YUMMER1.7 mouse melanoma model. Pigment Cell Melanoma Res 2017, 30, 428–435, doi:10.1111/pcmr.12591.

31. Malosse, C.; Henri, S. Isolation of Mouse Dendritic Cell Subsets and Macrophages from the Skin. Methods Mol Biol 2016, 1423, 129–137, doi:10.1007/978-1-4939-3606-9_9.

32. Etzerodt, A.; Tsalkitzi, K.; Maniecki, M.; Damsky, W.; Delfini, M.; Baudoin, E.; Moulin, M.; Bosenberg, M.; Graversen, J.H.; Auphan-Anezin, N.; et al. Specific targeting of CD163(+) TAMs mobilizes inflammatory monocytes and promotes T cell-mediated tumor regression. J Exp Med 2019, 216, 2394–2411, doi:10.1084/jem.20182124.

33. Muller, S.; Kohanbash, G.; Liu, S.J.; Alvarado, B.; Carrera, D.; Bhaduri, A.; Watchmaker, P.B.; Yagnik, G.; Di Lullo, E.; Malatesta, M.; et al. Single-cell profiling of human gliomas reveals macrophage ontogeny as a basis for regional differences in macrophage activation in the tumor microenvironment. Genome Biol 2017, 18, 234, doi:10.1186/s13059-017-1362-4.

34. Casanova, M.L.; Larcher, F.; Casanova, B.; Murillas, R.; Fernandez-Acenero, M.J.; Villanueva, C.; Martinez-Palacio, J.; Ullrich, A.; Conti, C.J.; Jorcano, J.L. A critical role for ras-mediated, epidermal growth factor receptor-dependent angiogenesis in mouse skin carcinogenesis. Cancer Res 2002, 62, 3402–3407.

35. Opzoomer, J.W.; Anstee, J.E.; Dean, I.; Hill, E.J.; Bouybayoune, I.; Caron, J.; Muliaditan, T.; Gordon, P.; Sosnowska, D.; Nuamah, R.; et al. Macrophages orchestrate the expansion of a proangiogenic perivascular niche during cancer progression. Sci Adv 2021, 7, eabg9518, doi:10.1126/sciadv.abg9518.

36. Summers, K.M.; Bush, S.J.; Hume, D.A. Network analysis of transcriptomic diversity amongst resident tissue macrophages and dendritic cells in the mouse mononuclear phagocyte system. PLoS Biol 2020, 18, e3000859, doi:10.1371/journal.pbio.3000859.

37. Casanova-Acebes, M.; Dalla, E.; Leader, A.M.; LeBerichel, J.; Nikolic, J.; Morales, B.M.; Brown, M.; Chang, C.; Troncoso, L.; Chen, S.T.; et al. Tissue-resident macrophages provide a pro-tumorigenic niche to early NSCLC cells. Nature 2021, 595, 578–584, doi:10.1038/s41586-021-03651-8.

38. Wang, P.L.; Yim, A.K.Y.; Kim, K.W.; Avey, D.; Czepielewski, R.S.; Colonna, M.; Milbrandt, J.; Randolph, G.J. Peripheral nerve resident macrophages share tissue-specific programming and features of activated microglia. Nat Commun 2020, 11, 2552, doi:10.1038/s41467-020-16355-w.

39. Nirmal, A.J.; Maliga, Z.; Vallius, T.; Quattrochi, B.; Chen, A.A.; Jacobson, C.A.; Pelletier, R.J.; Yapp, C.; Arias-Camison, R.; Chen, Y.A.; et al. The Spatial Landscape of Progression and Immunoediting in Primary Melanoma at Single-Cell Resolution. Cancer Discov 2022, 12, 1518–1541, doi:10.1158/2159-8290.CD-21-1357.

40. Martinek, J.; Lin, J.; Kim, K.I.; Wang, V.G.; Wu, T.C.; Chiorazzi, M.; Boruchov, H.; Gulati, A.; Seeniraj, S.; Sun, L.; et al. Transcriptional profiling of macrophages in situ in metastatic melanoma reveals localization-dependent phenotypes and function. Cell Rep Med 2022, 3, 100621, doi:10.1016/j.xcrm.2022.100621.

41. Schoenfeld, D.A.; Merkin, R.D.; Moutafi, M.; Martinez, S.; Adeniran, A.; Kumar, D.; Jilaveanu, L.; Hurwitz, M.; Rimm, D.L.; Kluger, H.M. Location matters: LAG3 levels are lower in renal cell carcinoma metastatic sites compared to primary tumors, and expression at metastatic sites only may have prognostic importance. Front Oncol 2022, 12, 990367, doi:10.3389/fonc.2022.990367.

42. Kielbassa, K.; Vegna, S.; Ramirez, C.; Akkari, L. Understanding the Origin and Diversity of Macrophages to Tailor Their Targeting in Solid Cancers. Front Immunol 2019, 10, 2215, doi:10.3389/fimmu.2019.02215.

43. Eisinger, S.; Sarhan, D.; Boura, V.F.; Ibarlucea-Benitez, I.; Tyystjarvi, S.; Oliynyk, G.; Arsenian-Henriksson, M.; Lane, D.; Wikstrom, S.L.; Kiessling, R.; et al. Targeting a scavenger receptor on tumor-associated macrophages activates tumor cell killing by natural killer cells. Proc Natl Acad Sci U S A 2020, 117, 32005–32016, doi:10.1073/pnas.2015343117.

44. Adams, R.; Osborn, G.; Mukhia, B.; Laddach, R.; Willsmore, Z.; Chenoweth, A.; Geh, J.L.C.; MacKenzie Ross, A.D.; Healy, C.; Barber, L.; et al. Influencing tumor-associated macrophages in malignant melanoma with monoclonal antibodies. Oncoimmunology 2022, 11, 2127284, doi:10.1080/2162402X.2022.2127284.

45. Wang, S.; Liu, G.; Li, Y.; Pan, Y. Metabolic Reprogramming Induces Macrophage Polarization in the Tumor Microenvironment. Front Immunol 2022, 13, 840029, doi:10.3389/fimmu.2022.840029.

46. Liu, P.S.; Wang, H.; Li, X.; Chao, T.; Teav, T.; Christen, S.; Di Conza, G.; Cheng, W.C.; Chou, C.H.; Vavakova, M.; et al. alpha-ketoglutarate orchestrates macrophage activation through metabolic and epigenetic reprogramming. Nat Immunol 2017, 18, 985–994, doi:10.1038/ni.3796.

47. Kelly, B.; O’Neill, L.A. Metabolic reprogramming in macrophages and dendritic cells in innate immunity. Cell Res 2015, 25, 771–784, doi:10.1038/cr.2015.68.

48. Lin, J.D.; Nishi, H.; Poles, J.; Niu, X.; McCauley, C.; Rahman, K.; Brown, E.J.; Yeung, S.T.; Vozhilla, N.; Weinstock, ; et al. Single-cell analysis of fate-mapped macrophages reveals heterogeneity, including stem-like properties, during atherosclerosis progression and regression. JCI Insight 2019, 4, doi:10.1172/jci.insight.124574.

49. Chakarov, S.; Lim, H.Y.; Tan, L.; Lim, S.Y.; See, P.; Lum, J.; Zhang, X.M.; Foo, S.; Nakamizo, S.; Duan, K.; et al. Two distinct interstitial macrophage populations coexist across tissues in specific subtissular niches. Science 2019, 363, pdoi:10.1126/science.aau0964.

50. Mittal, A.; Wang, M.; Vidyarthi, A.; Yanez, D.; Pizzurro, G.; Thakral, D.; Tracy, E.; Colegio, O.R. Topical arginase inhibition decreases growth of cutaneous squamous cell carcinoma. Sci Rep 2021, 11, 10731, doi:10.1038/s41598-021-90200-y.

