## Supplementary Material for "Functionally and metabolically divergent melanoma-associated macrophages originate from common bone-marrow precursors"

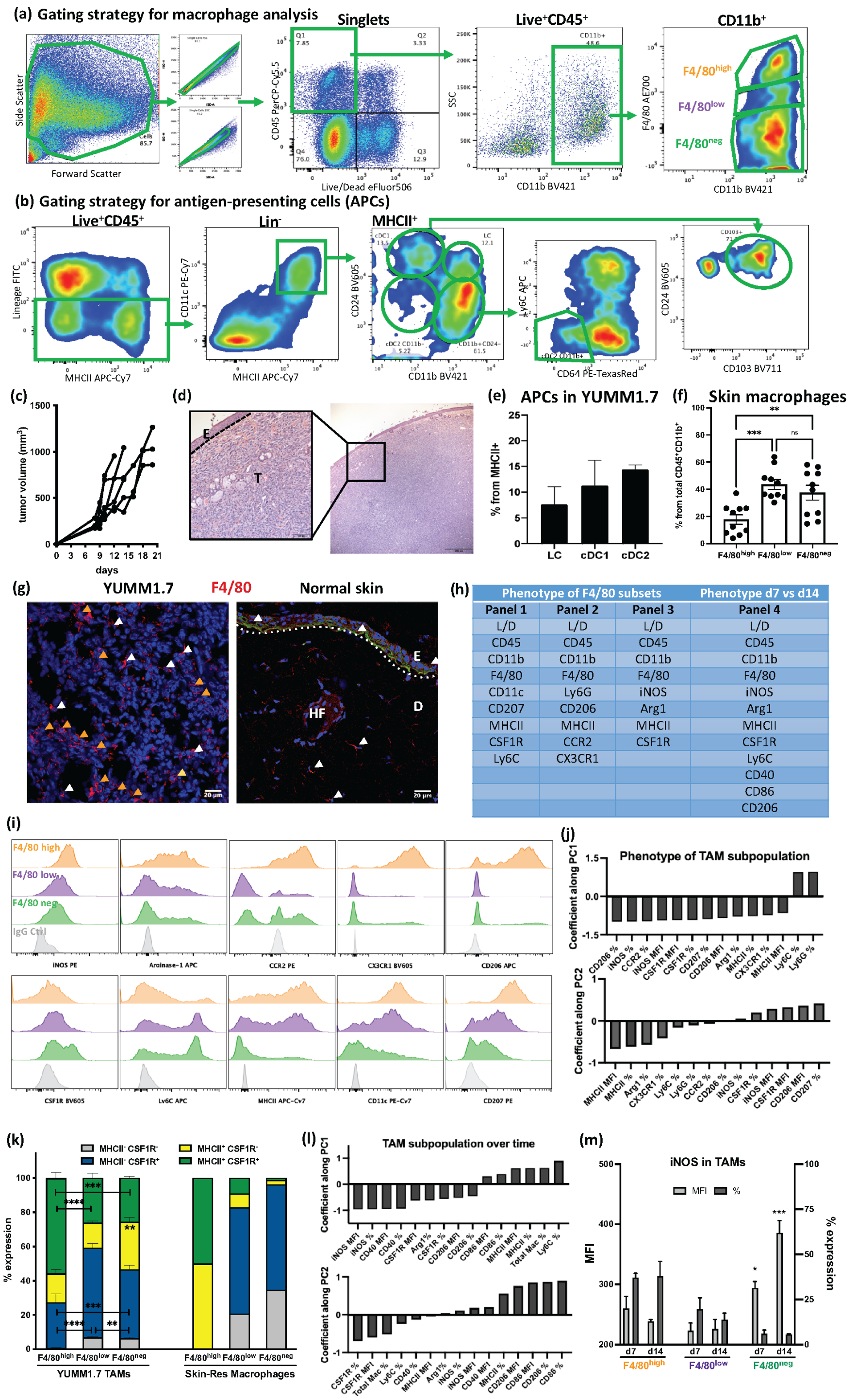


| Phenotype of F4/80 subsets | | | Phenotype d7 vs d14 |
|---|---|---|---|
| Panel 1 | Panel 2 | Panel 3 | Panel 4 |
| L/D | L/D | L/D | L/D |
| CD45 | CD45 | CD45 | CD45 |
| CD11b | CD11b | CD11b | CD11b |
| F4/80 | F4/80 | F4/80 | F4/80 |
| CD11c | Ly6G | iNOS | iNOS |
| CD207 | CD206 | Arg1 | Arg1 |
| MHCII | MHCII | MHCII | MHCII |
| CSF1R | CCR2 | CSF1R | CSF1R |
| Ly6C | CX3CR1 | | Ly6C |
| | | | CD40 |
| | | | CD86 |
| | | | CD206 |

**Supplementary Figure 1. Tumor-associated macrophage subsets in the i.d.-injected YUMM mouse melanoma model.** **(a)** Flow cytometry gating strategy for analyzing F4/80 macrophage subpopulations in YUMM1.7 tumors. **(b)** Flow cytometry gating strategy for determining APC subsets in YUMM1.7 tumors. **(c)** Individual tumor growth curves of the YUMM1.7 cell line in C57Bl/6J. **(d)** H&E and detail from i.d. YUMM1.7 tumor at day 14, injected in Myb mice. E = epidermis, T = tumor. **(e)** FACS quantification of skin myeloid APC subsets in YUMM1.7 tumors in Myb mice. n=4. **(f)** Skin-resident macrophages determined by their F4/80 expression in C57Bl/6J. n=10, pooled from at least 3 independent experiments. **(g)** Immunofluorescent staining for F4/80$^+$ TAMs infiltration in YUMM1.7 melanoma tumor and adjacent dermis 'D' and epidermis 'E'. White triangles indicate single F4/80$^+$ cells, while orange triangles point out clusters of TAMs. **(h)** Marker panels used for phenotyping myeloid cells infiltrating YUMM1.7 tumors and normal skin. **(i)** Representative histograms of flow cytometry marker expression analyzed in the TAM subset comparisons. **(j)** Coefficients along PC1 (top) and PC2 (bottom) for the PCA of phenotypic markers of F4/80 subpopulations in tumor vs skin. As indicated, either percentage of marker expression (%) or % and MFI were included in the analysis. **(k)** Analysis of monocyte-to-macrophage marker co-expression (CSF1R and MHCII) in TAM subpopulations from YUMM1.7 tumors, and a control skin. Statistical analysis showing comparisons between marker-expressing subsets within TAMs. **(l)** Coefficients along PC1 (top) and PC2 (bottom) for the PCA of phenotypic markers of F4/80 subpopulations in tumors day 7 vs day 14. As indicated, either percentage of marker expression (%) or % and MFI were included in the analysis. **(m)** Detail of the evolution of percentage and MFI of M1-marker iNOS expression in F4/80 TAM subsets. n=5 mice per TAM group, pooled from 3 independent experiments. ns= not significant, *p<0.05, **p<0.01, ***p<0.001.

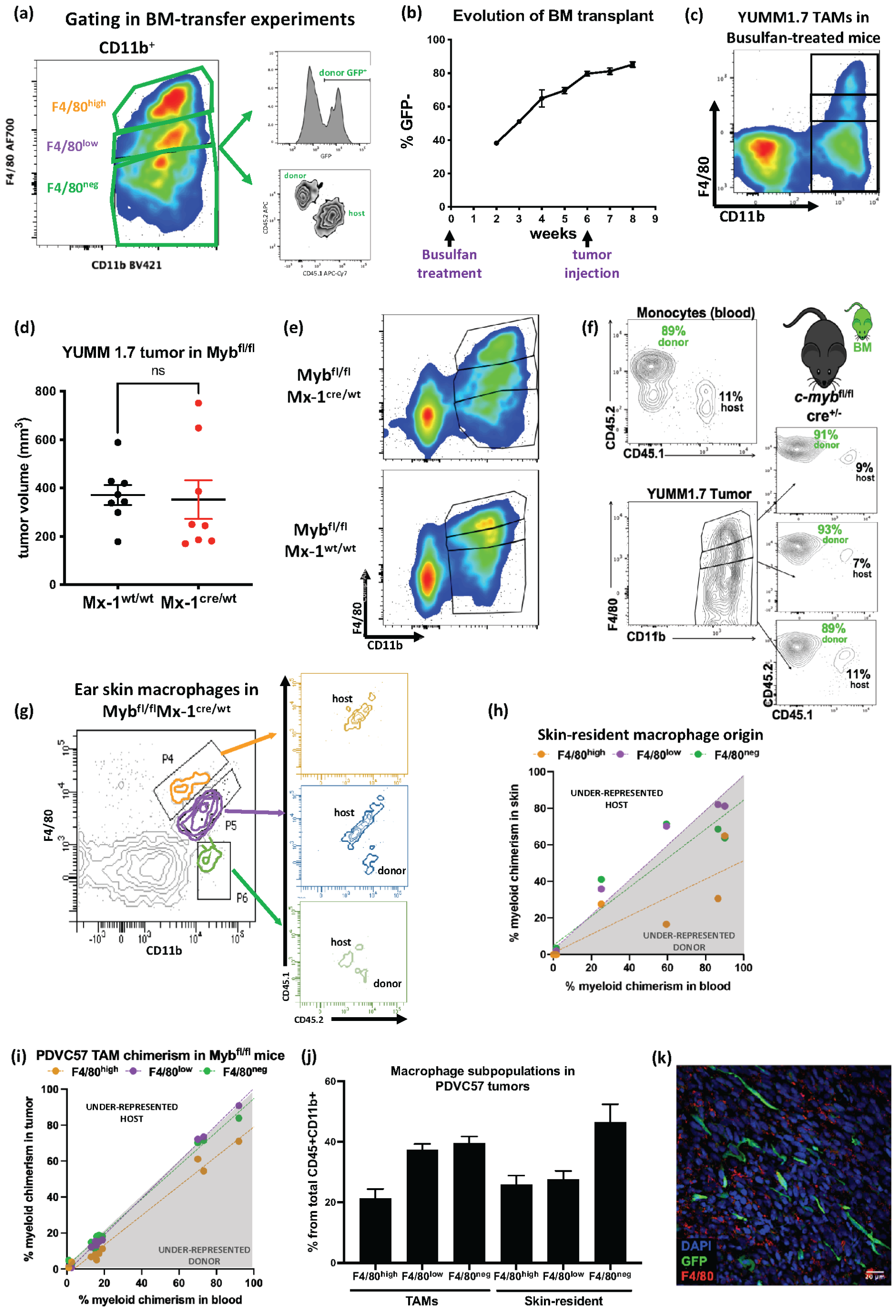

(a)
Gating in BM-transfer experiments
CD11b+
F4/80high
F4/80low
F4/80neg
F4/80 AF700
CD11b BV421
donor GFP+
donor
host
(b)
Evolution of BM transplant
% GFP-
weeks
Busulfan treatment
tumor injection
(c)
YUMM1.7 TAMs in Busulfan-treated mice
F4/80
CD11b
(d)
YUMM 1.7 tumor in Mybfl/fl
ns
tumor volume (mm3)
Mx-1wt/wt
Mx-1cre/wt
(e)
Mybfl/fl Mx-1cre/wt
Mybfl/fl Mx-1wt/wt
F4/80
CD11b
(f)
Monocytes (blood)
89% donor
11% host
CD45.2
CD45.1
BM
c-mybfl/fl cre+/-
YUMM1.7 Tumor
91% donor
9% host
93% donor
7% host
89% donor
11% host
F4/80
CD11b
(g)
Ear skin macrophages in Mybfl/flMx-1cre/wt
P4
P5
P6
F4/80
CD11b
host
host
donor
host
donor
CD45.1
CD45.2
(h)
Skin-resident macrophage origin
F4/80high F4/80low F4/80neg
UNDER-REPRESENTED HOST
UNDER-REPRESENTED DONOR
% myeloid chimerism in skin
% myeloid chimerism in blood
(i)
PDVC57 TAM chimerism in Mybfl/fl mice
F4/80high F4/80low F4/80neg
UNDER-REPRESENTED HOST
UNDER-REPRESENTED DONOR
% myeloid chimerism in tumor
% myeloid chimerism in blood
(j)
Macrophage subpopulations in PDVC57 tumors
% from total CD45+CD11b+
F4/80high F4/80low F4/80neg F4/80high F4/80low F4/80neg
TAMs
Skin-resident
(k)
DAPI
GFP
F4/80

**Supplementary Figure 2. Origin of melanoma-associated macrophage subsets. (a)** Flow cytometry strategy for determining TAM origin in YUMM1.7 tumors. Fate-mapping of BM-derived cells was identified by CD45.1/CD45.2 expression and validated by $GFP^{+}$. **(b)** Assessment of BM engraftment after Busulfan treatment and transplant. Blood samples were extracted and analyzed weekly. The graph shows the percentage of $GFP^{-}$ cells from total $CD45^{+}$ cells in $GFP^{+}$ mice. n=5. **(c)** Representative plot of TAM subpopulations in Busulfan-treated mice 8 weeks after transplant. **(d)** YUMM1.7 tumor size in $Myb^{fl/fl}$ mice at day 14, at the time of processing for analysis. **(e)** Representative flow cytometry plots of TAM subpopulations in YUMM1.7 tumors in $Myb^{fl/fl}Mx\text{-}1^{cre/wt}$ and $Myb^{fl/fl}Mx\text{-}1^{wt/wt}$ mice. **(f)** Schematic of the analysis of macrophage origin in tumor samples in $Myb^{fl/fl}Mx\text{-}1^{cre/wt}$ mice, and comparison with blood monocyte chimerism. Host immune cells are $CD45.1^{+}GFP^{-}$ and donor cells are $CD45.2^{+}$ and $GFP^{+}$. **(g)** Representative plots of the analysis of the origin of macrophages in ear skin from BM-transplanted Myb mice. **(h)** Assessment of macrophage origin in skin of tumor-bearing $Myb^{fl/fl}Mx\text{-}1^{cre/wt}$ and $Myb^{fl/fl}Mx\text{-}1^{wt/wt}$mice. Linear regressions were performed to compare the chimerism of myeloid cells observed in circulation and the chimerism of myeloid tumor-infiltrating cells. Skin samples, n=8, pooled from 2 independent experiments. **(i)** Assessment of macrophage origin in PDVC57 tumors in $Myb^{fl/fl}Mx\text{-}1^{cre/wt}$ and $Myb^{fl/fl}Mx\text{-}1^{wt/wt}$ mice. Linear regressions were performed to compare the chimerism of myeloid cells observed in circulation and the chimerism of myeloid tumor-infiltrating cells. TAM subsets were analyzed separately. Tumor samples, n=10, pooled from 2 independent experiments. **(j)** Quantification of TAMs and skin-resident macrophage subsets based on F4/80 expression in i.d.-injected PDVC57 tumors in C57Bl/6J. n=6-10, pooled from 2 independent experiments. **(k)** Immunofluorescence staining on YUMM1.7 tumor from Busulfan-treated mice, showing only $GFP^{+}$ stromal cells.

(a)

| | | |
|---|---|---|
| Sample_1 | F4/80high | 161416 |
| | F4/80low | 467320 |
| Sample_2 | F4/80high | 522565 |
| | F4/80low | 1471916 |
| Sample_3 | F4/80high | 218228 |
| | F4/80low | 624604 |

(b)

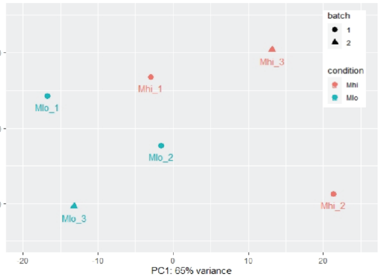


(c)

F4/80high TAMs

| | Log2FoldChange | p-value (adj) |
|---|---|---|
| Gm28049 | 6.584604715 | 0.009240763 |
| St18 | 5.615315739 | 0.047023634 |
| Gbgt1 | 3.83766266 | 0.000686275 |
| Mrvi1 | 3.799213555 | 0.022372385 |
| Gm6710 | 3.400979485 | 0.042578028 |
| RP23-26P10.1 | 3.304597769 | 0.00095194 |
| Cd163 | 3.152898245 | 0.02167177 |
| Slco2b1 | 3.004562616 | 0.005421037 |
| Rapgef5 | 2.936676966 | 0.023225688 |
| Gm15880 | 2.810271781 | 0.022372385 |
| Ms4a14 | 2.621480237 | 0.000211194 |
| Scamp5 | 2.468269971 | 0.000584185 |
| Spin4 | 2.398096625 | 0.029308747 |
| Zdhhc14 | 2.387773517 | 0.014463673 |
| Hpgds | 2.372577924 | 0.004765114 |
| Lonrf3 | 2.305200623 | 0.020415803 |
| Apold1 | 2.233013864 | 0.029308747 |
| Mrc1 | 2.219736182 | 0.010974313 |
| Itga9 | 2.208618885 | 0.005793151 |
| Igf1 | 2.163549432 | 0.007743335 |
| Gm10134 | 2.152394047 | 0.001137803 |
| Tdrkh | 2.017959613 | 0.010629957 |
| C1qa | 1.960986541 | 0.047382687 |
| Gpr34 | 1.948690168 | 0.005307293 |
| Ms4a7 | 1.935309663 | 0.024302447 |
| Arhgap19 | 1.931112894 | 0.00145479 |
| Cadm1 | 1.907844223 | 0.011228814 |
| Olfml3 | 1.899902242 | 0.002047071 |
| Stab1 | 1.832088096 | 0.034524385 |
| C4b | 1.772663925 | 0.01413023 |
| Blnk | 1.770030187 | 0.021973892 |
| Prune2 | 1.704464226 | 0.020415803 |
| Mertk | 1.665602583 | 0.022509876 |
| Gdpd5 | 1.646956437 | 0.022509876 |
| Gm37795 | 1.618343355 | 0.030222198 |
| Bbs7 | 1.616514749 | 0.036947432 |
| Abcc3 | 1.61598911 | 0.038339054 |
| Ccl8 | 1.601805093 | 0.030222198 |
| Frmd4b | 1.57734899 | 0.000506847 |
| Rab3il1 | 1.545887596 | 0.035665271 |
| Ctla2b | 1.503755412 | 0.019820169 |
| Cd38 | 1.468646345 | 0.027659319 |
| RP24-180F7.2 | 1.46376072 | 0.014963899 |
| Sepp1 | 1.458027668 | 0.02395971 |
| Abcb1b | 1.361607969 | 0.001137803 |
| Zmynd15 | 1.302103388 | 0.049109972 |
| Pltp | 1.278872876 | 0.041641002 |
| Pdgfc | 1.214844511 | 0.042578028 |
| Cebpa | 1.1662205 | 0.012421469 |
| Maml2 | 1.078892543 | 0.022372385 |
| Cd72 | 1.055101448 | 0.042578028 |

F4/80low TAMs

| | log2FoldChange | p-value (adj) |
|---|---|---|
| Sytl4 | -7.847994864 | 0.000502805 |
| Grm1 | -7.463531415 | 0.007658365 |
| Afap1l2 | -7.340588703 | 0.000584185 |
| Sh3tc2 | -6.159471225 | 0.013968525 |
| Gm37334 | -5.309857158 | 4.75E-05 |
| Abca6 | -5.239063108 | 0.00572596 |
| Cubn | -5.204547692 | 0.011228814 |
| 6230415J03Rik | -5.180191644 | 0.004088306 |
| Gm37873 | -5.126793775 | 0.023225688 |
| Cttnbp2 | -5.07996975 | 4.75E-05 |
| AW121686 | -4.920024497 | 0.022372385 |
| Gm37115 | -4.830402755 | 0.000146079 |
| Epha4 | -4.787700318 | 0.002253627 |
| BC039771 | -3.94560155 | 0.023687276 |
| Grm6 | -3.867535917 | 0.008313685 |
| Btnl9 | -3.800560949 | 0.032456106 |
| Ccdc64b | -3.725221462 | 0.004369468 |
| Ttc26 | -3.636662517 | 0.002592003 |
| Arhgef28 | -3.610244951 | 0.004918731 |
| Gm10282 | -3.418129862 | 0.047382687 |
| Ighg1 | -3.244328781 | 0.000130998 |
| Gm7292 | -2.922087549 | 0.032008481 |
| Dcstamp | -2.750728894 | 0.002592003 |
| Gm29666 | -2.744594656 | 0.047398534 |
| Bmx | -2.427075833 | 0.039614528 |
| Ctcflos | -2.424551074 | 0.001616943 |
| Mcmdc2 | -2.38201597 | 0.048562434 |
| Gm15513 | -2.378260731 | 0.018447742 |
| Cldn15 | -2.374790715 | 0.042635601 |
| Pbld2 | -2.274623258 | 0.017163329 |
| Gm37760 | -2.15590194 | 0.045151748 |
| Smpd3 | -2.138543983 | 0.000584185 |
| Abcc2 | -2.063266298 | 0.012898923 |
| Gm28187 | -2.044480862 | 0.009307088 |
| Dach1 | -1.962524055 | 0.023687276 |
| Ffar2 | -1.96203876 | 0.004918731 |
| Zbtb16 | -1.931605511 | 0.026998616 |
| Nxpe4 | -1.917520391 | 6.43E-06 |
| I830127L07Rik | -1.884296572 | 0.002253627 |
| Prg2 | -1.869431428 | 0.001137803 |
| Slco4c1 | -1.81998291 | 0.00095194 |
| Chil3 | -1.806328695 | 0.000502805 |
| Sag | -1.751140706 | 0.047023634 |
| Vcan | -1.751118976 | 0.018851697 |
| Mylk3 | -1.732723562 | 0.047382687 |
| Ear2 | -1.718321127 | 0.025561278 |
| Gata1 | -1.662998314 | 0.036938585 |
| Cd177 | -1.637362825 | 0.024302447 |
| Ccr3 | -1.635771299 | 0.006122297 |
| Ifitm6 | -1.627062142 | 0.002253627 |
| Col27a1 | -1.618677663 | 0.040805213 |

(d)

F4/80high TAMs

| NAME | SIZE | NES | NOM p-val | FDR q-val |
|---|---|---|---|---|
| HALLMARK_E2F_TARGETS | 192 | 2.464 | 0.000 | 0.000 |
| HALLMARK_MYC_TARGETS_V1 | 191 | 2.448 | 0.000 | 0.000 |
| HALLMARK_MYC_TARGETS_V2 | 58 | 2.218 | 0.000 | 0.000 |
| HALLMARK_G2M_CHECKPOINT | 182 | 1.772 | 0.000 | 0.003 |
| HALLMARK_OXIDATIVE_PHOSPHORYLATION | 194 | 1.551 | 0.000 | 0.020 |
| HALLMARK_PROTEIN_SECRETION | 91 | 1.290 | 0.074 | 0.175 |
| HALLMARK_ALLOGRAFT_REJECTION | 185 | 1.285 | 0.058 | 0.159 |
| HALLMARK_PEROXISOME | 92 | 1.250 | 0.090 | 0.182 |

F4/80low TAMs

| NAME | SIZE | NES | NOM p-val | FDR q-val |
|---|---|---|---|---|
| HALLMARK_INTERFERON_ALPHA_RESPONSE | 90 | 2.146 | 0.000 | 0.000 |
| HALLMARK_INTERFERON_GAMMA_RESPONSE | 183 | 2.096 | 0.000 | 0.000 |
| HALLMARK_TNFA_SIGNALING_VIA_NFKB | 191 | 2.067 | 0.000 | 0.000 |
| HALLMARK_EPITHELIAL_MESENCHYMAL_TRANSITION | 180 | 1.958 | 0.000 | 0.000 |
| HALLMARK_HYPOXIA | 178 | 1.881 | 0.000 | 0.001 |
| HALLMARK_COAGULATION | 104 | 1.847 | 0.000 | 0.001 |
| HALLMARK_INFLAMMATORY_RESPONSE | 185 | 1.776 | 0.000 | 0.004 |
| HALLMARK_UV_RESPONSE_DN | 137 | 1.714 | 0.000 | 0.006 |
| HALLMARK_REACTIVE_OXIGEN_SPECIES_PATHWAY | 46 | 1.696 | 0.007 | 0.006 |
| HALLMARK_APOPTOSIS | 156 | 1.666 | 0.000 | 0.006 |
| HALLMARK_APICAL_JUNCTION | 166 | 1.578 | 0.002 | 0.012 |
| HALLMARK_MYOGENESIS | 163 | 1.562 | 0.000 | 0.013 |
| HALLMARK_ESTROGEN_RESPONSE_LATE | 163 | 1.561 | 0.000 | 0.012 |
| HALLMARK_ANGIOGENESIS | 34 | 1.481 | 0.028 | 0.026 |
| HALLMARK_ESTROGEN_RESPONSE_EARLY | 171 | 1.379 | 0.013 | 0.067 |
| HALLMARK_KRAS_SIGNALING_UP | 176 | 1.376 | 0.011 | 0.064 |
| HALLMARK_COMPLEMENT | 171 | 1.351 | 0.026 | 0.076 |

(e)

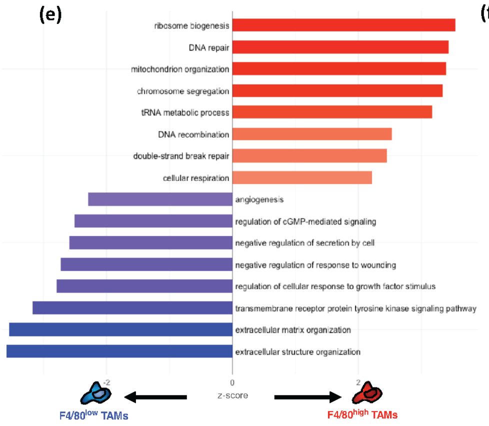


(f)

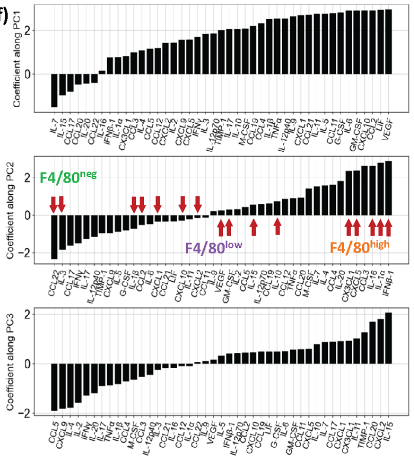


(g)

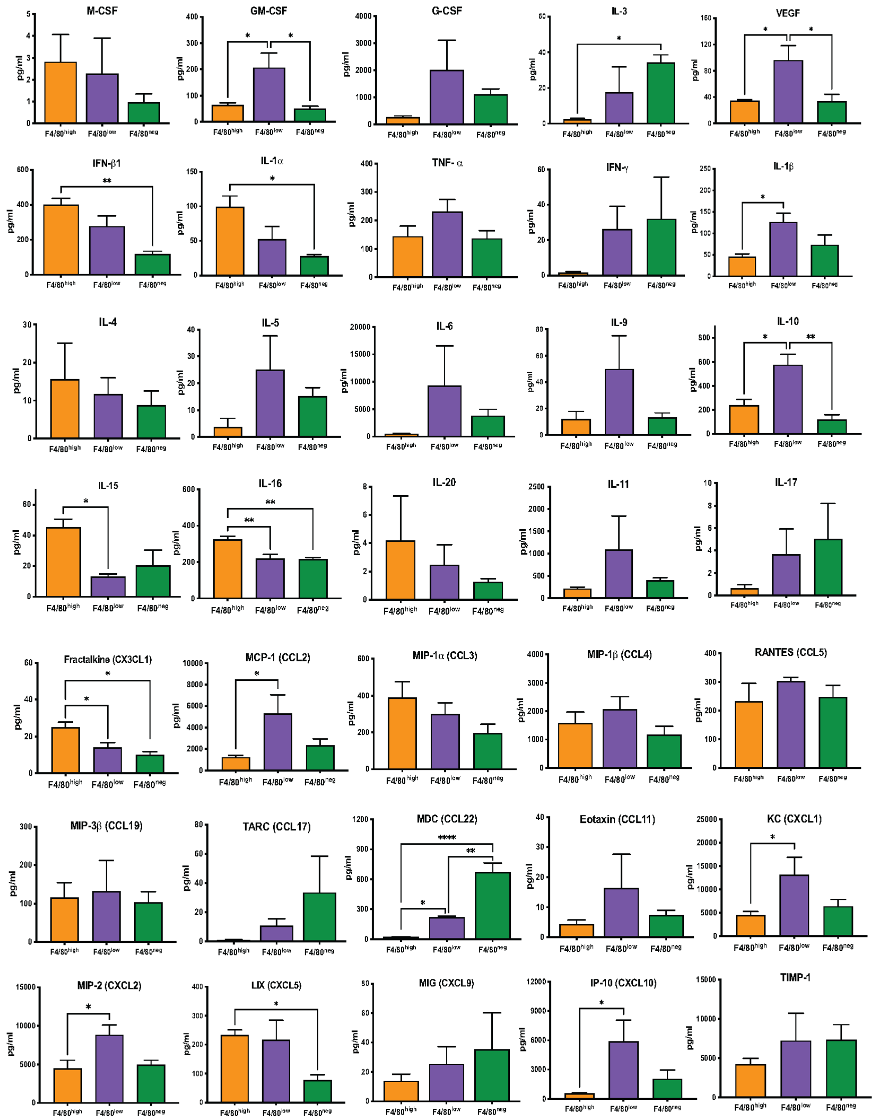

**Supplementary Figure 3. Gene expression profiling and functional analysis of TAM subsets in YUMM1.7 tumors**. **(a)** Total number of cells sorted from TAM subsets for bulk RNAseq. **(b)** PCA embedding of bulk RNAseq samples after batch-correction. **(c)** Top 50 differentially-expressed genes (DEGs) for each F4/80 subset (p-adj < 0.05, abs(log2FC) > 1). **(d)** GSEA results showing significantly upregulated Hallmark gene sets from the mouse MSigDB, with a FDR<25% for each TAM subset. **(e)** Enrichment of F4/80$^{high}$ (right, red) and F4/80$^{low}$ (left, blue) DEGs (protein-coding genes only) for gene ontology (GO) biological process terms. Bar lengths represent associated z-score. **(f)** Coefficients along PC1 (top), PC2 (middle), and PC3 (bottom) for the PCA of multiplex protein secretion assay performed on sorted F4/80$^{high}$, F4/80$^{low}$ and F4/80$^{neg}$ subsets from YUMM1.7 samples. Along PC2, samples separated and clustered by F4/80 expression level. Red arrows indicate the proteins that showed significantly different levels of secretion, detailed in (g). **(g)** Individual bar graphs of protein secretion from sorted TAM subsets which showed significant differences or trends between them, with relevance to immune function in the TME. *p<0.05, **p<0.01, ****p<0.0001.
